## Supplementary material for "Structure and rational engineering of the PglX methyltransferase and specificity factor for BREX phage defence": Merged Supplemental

[b]New England Biolabs, 240 County Road, Ipswich, MA 01938, USA.

[c]Faculty of Health and Life Sciences, Northumbria University, Newcastle Upon Tyne, NE1 8ST, UK.

[d]Institute of Infection, Veterinary and Ecological Sciences, University of Liverpool, Liverpool, L69 7ZB, UK.

Keywords: BREX, phage defence, PglX, methyltransferase, Ocr

### Supplementary Figures

#### Supplementary Figure S1

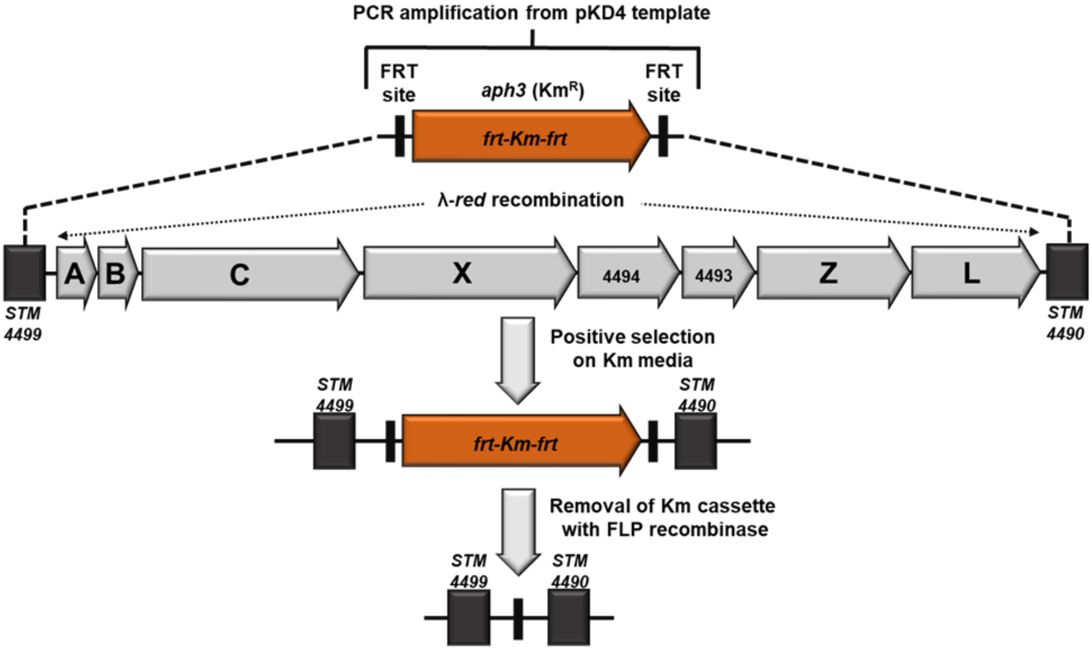


**Supplementary Figure S1. Generation of *S*. Typhimurium D23580 BREX phage defence island knockout using Lambda red recombination**. All genetic material between *STM4498* and *STM4491* (inclusive) was removed, and the kanamycin resistance cassette was cured.

### Supplementary Figure S2

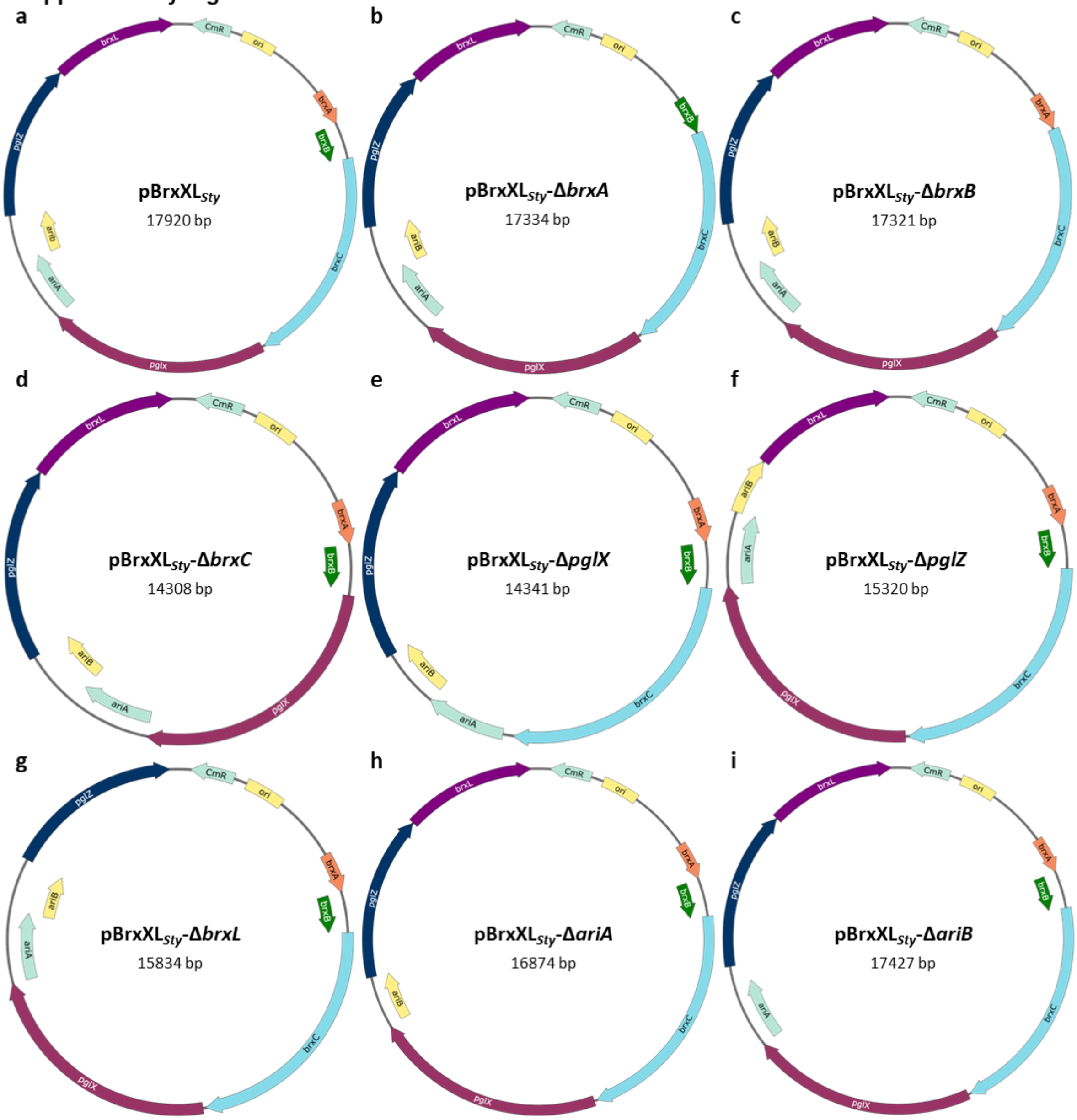


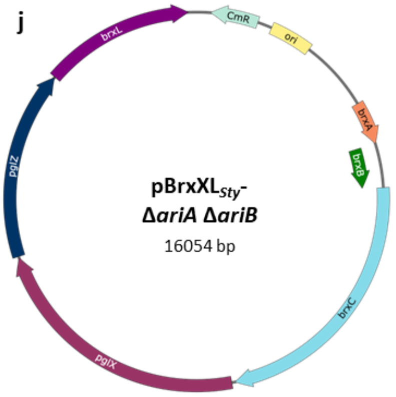


**Supplementary Figure S2.** ***Salmonella*** **D23580 BREX plasmids produced for this study**. The initial pGGA-based pBrxXL$_{Sty}$ plasmid (**a**) was previously produced by golden gate assembly[33] and used as a template for the production of gene knockout derivatives (**b-j**) by Gibson assembly.

### Supplementary Figure S3

| Phage | BREX sites | pBrxXL$_{Sty}$ | pBrxXL$_{Sty}$-Δ*brxL* | pBrxXL$_{Sty}$-Δ*ariA*Δ*ariB* |
|---|---|---|---|---|
| TB34 | 120 | 0.03 ± 0.02 | 7.50 x10$^{-6}$ ± 3.46 x10$^{-6}$ | 0.01 ± 2.96 x10$^{-3}$ |
| Trib | 0 | 1.19 ± 0.11 | 0.70 ± 0.27 | 0.94 ± 0.23 |
| Baz | 0 | 1.35 ± 0.46 | 1.34 ± 0.81 | 2.08 ± 1.67 |
| Alma | 110 | 0.03 ± 3.21 x10$^{-3}$ | 2.7 x10$^{-4}$ ± 1.61 x10$^{-4}$ | 6.35 x 10$^{-3}$ ± 4.39 x10$^{-3}$ |
| Pau | 84 | 0.19 ± 0.13 | 3.73 x10$^{-4}$ ± 2.65 x 10$^{-4}$ | 0.14 ± 0.27 |
| BB1 | 51 | 0.11 ± 0.05 | 0.27 ± 0.91 | <4.29 x10$^{-8}$ ± 1.61 x10$^{-8}$ |
| Jura | 0 | 2.58 ± 0.11 | 1.95 ± 1.37 | 1.82 ± 0.53 |
| Mak | 1 | 0.78 ± 0.10 | 3.77 ± 2.90 | 0.67 ± 0.12 |
| Bam | 1 | 1.06 ± 0.60 | 1.71 ± 0.69 | 0.59 ± 0.16 |
| CS16 | 13 | 0.03 ± 0.02 | 0.01 ± 2.39 x10$^{-3}$ | 0.06 ± 0.02 |
| Mav | 13 | 0.15 ± 0.02 | 0.06 ± 4.08 x10$^{-2}$ | 0.09 ± 0.10 |
| Sip | 84 | 0.04 ± 0.01 | 2.19 x10$^{-5}$ ± 2.57 x10$^{-4}$ | 5.81 x10$^{-3}$ ± 2.06 x10$^{-2}$ |

EOP ≥ 10$^{-1}$ | 10$^{-1}$ > EOP ≥ 10$^{-2}$ | 10$^{-2}$ > EOP ≥ 10$^{-3}$ | 10$^{-3}$ > EOP ≥ 10$^{-4}$ | 10$^{-4}$ > EOP

**Supplementary Figure S3. Wild type and mutant pBrxXL$_{Sty}$ plasmids show diverse defence against Durham collection phages**. EOPs of phages were tested against *E. coli* DH5α pBrxXL$_{Sty}$ plasmids with *E. coli* DH5α pTRB507 as control. Values are mean EOPs from triplicate data, shown with standard deviation.

**Supplementary Figure S4**

**a**

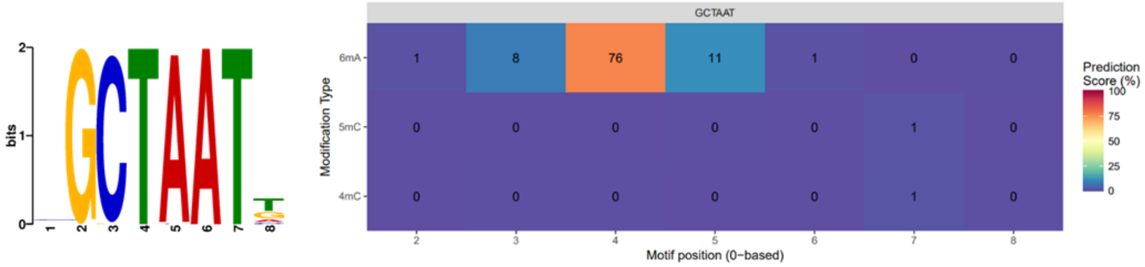


**b**

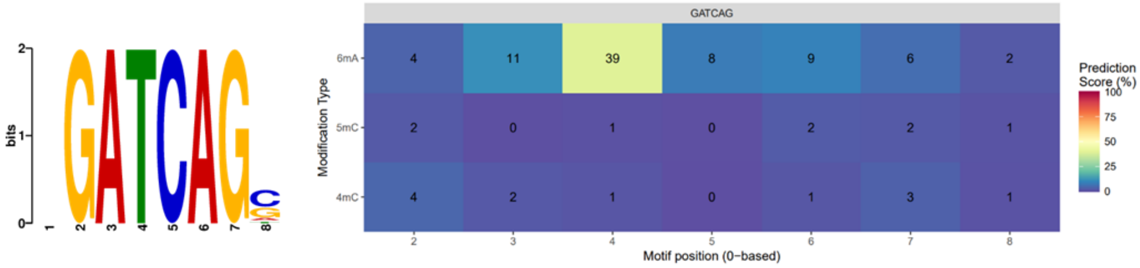


**Supplementary Figure S4. Methylation analysis detected BREX motifs**. Tombo (left) and nanodisco (right) analysis of genomic methylation from (**a**) *Escherichia fergusonii* and (**b**) *Salmonella* D23580 BREX systems. Tombo results represent enriched motifs around detected methylation signals, as identified by MEME (https://meme-suite.org/meme/). Nanodisco results show the type and position of DNA modification within DNA motifs identified by Tombo.

**Supplementary Figure S5**

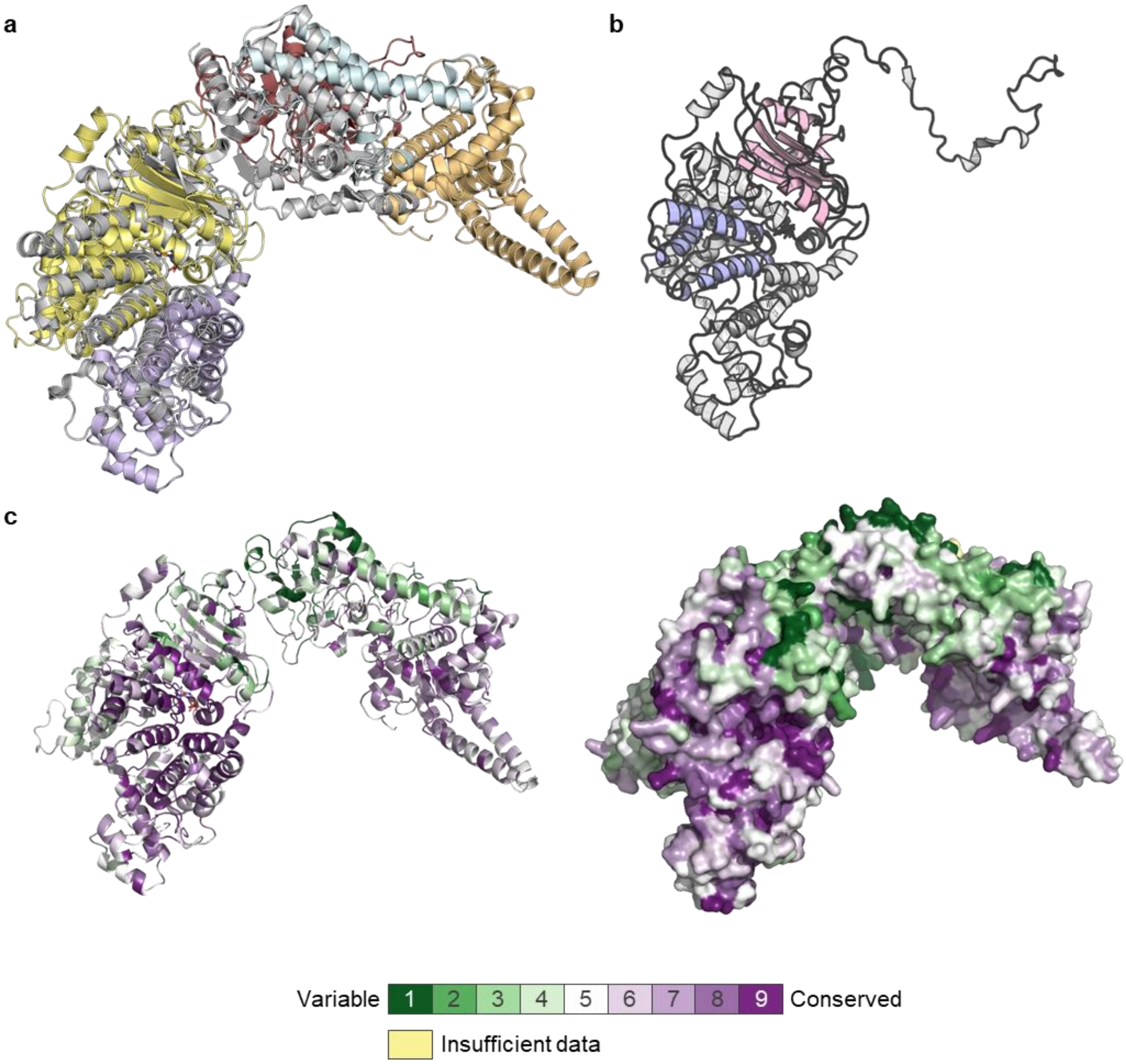


**Supplementary Figure S5. The methyltransferase and C-terminal regions of PglX are conserved.** (**a**) superposition of MmeI (5HR4) onto PglX (RMSD = 7.1 Å). (**b**) Position of the predicted MmeI-like DNA methyltransferase region (light blue) and adenine-specific DNA methylase region (light pink) within the N-terminal domain (gray) of PglX. (**c**) Ribbon and surface views of conserved residues in PglX, as calculated by ConSurf[43].

**Supplementary Figure S6**

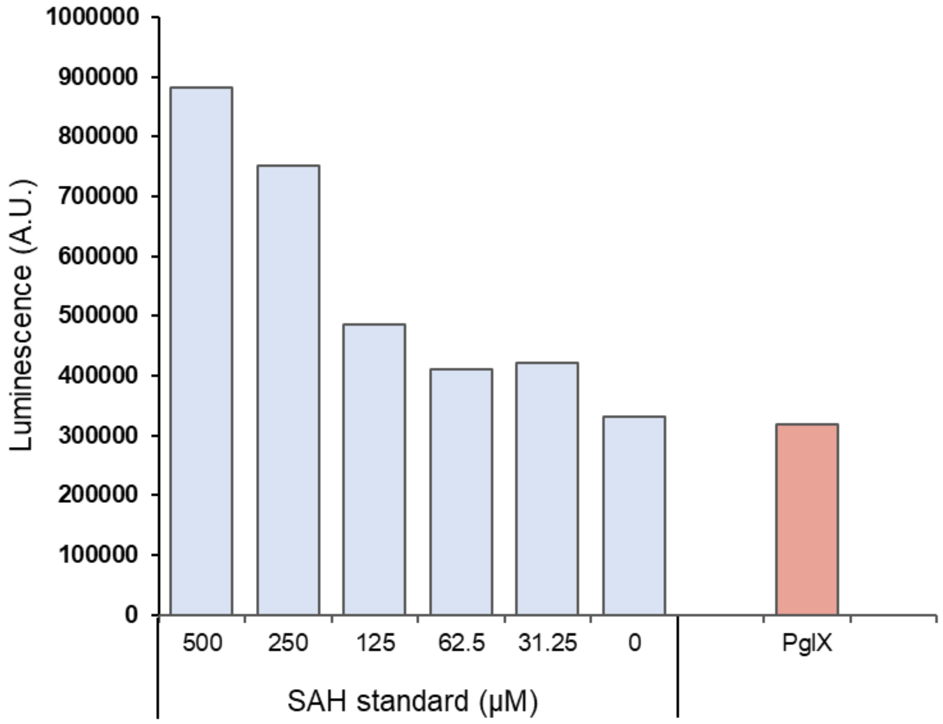


**Supplementary Figure S6. *In vitro* methyltransferase activity analysis of purified PglX shows no methyltransferase activity**. Activity of PglX relative to SAH positive controls measured indirectly through production of the SAH methylation reaction byproduct using the MTase-Glo kit (Promega). Data are representative of two replicates.

### Supplementary Figure S7

| Strain | EOP |
|---|---|
| pBrxXL$_{Sty}$-*ΔpglX* | 5.68 x10$^{-1}$ ± 3.94 x10$^{-1}$ |
| pBrxXL$_{Sty}$-*ΔpglX* pBAD30-*ocr* | 1.22 ± 0.8 |
| pBrxXL$_{Sty}$-*ΔpglX* pBAD30-*gp5* | 1.06 ± 0.35 |

**Supplementary Figure S7. Ocr and Gp5 do not activate the *Salmonella* PARIS system**. EOPs of TB34 tested against *E. coli* DH5α pBrxXL$_{Sty}$-*ΔpglX* co-expressing either Ocr or its *Salmonella* homologue, Gp5, with *E. coli* DH5α pTRB507 as control. Values are mean EOPs from triplicate data, shown with standard deviation.

### Supplementary Figure S8

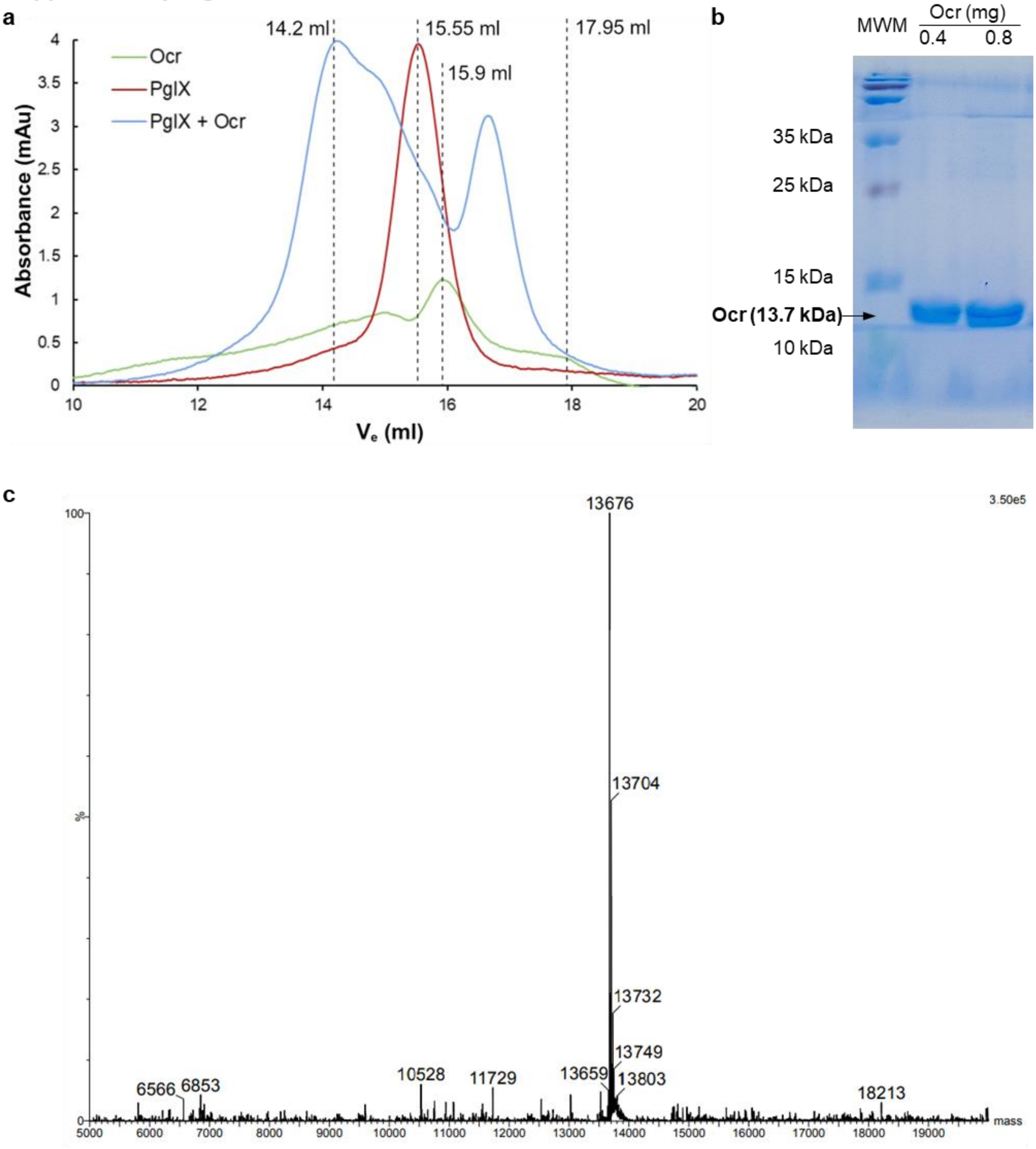


**Supplementary Figure S8. PglX directly interacts with Ocr *in vitro*.** (**a**) Analytical size exclusion chromatography of purified PglX and Ocr alongside PglX pre-incubated with Ocr. Elution volumes ($V_e$) of significant peaks are labelled. (**b**) SDS-PAGE analysis of purified Ocr. (**c**) Mass spectrometry results for purified Ocr.

**Supplementary Figure S9**

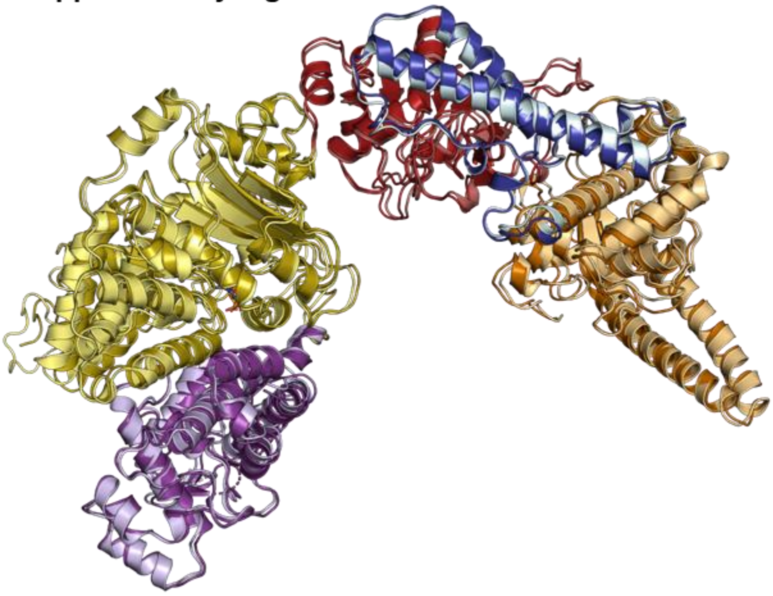

**Supplementary Figure S9.** Superposition of a protomer of PglX from the PglX-SAM structure (lighter shades), with PglX from the heterotetrameric PglX-SAM:Ocr complex (darker shades; RMSD = 1.34). Colored as per **Fig. 4**.

**Supplementary Figure S10**

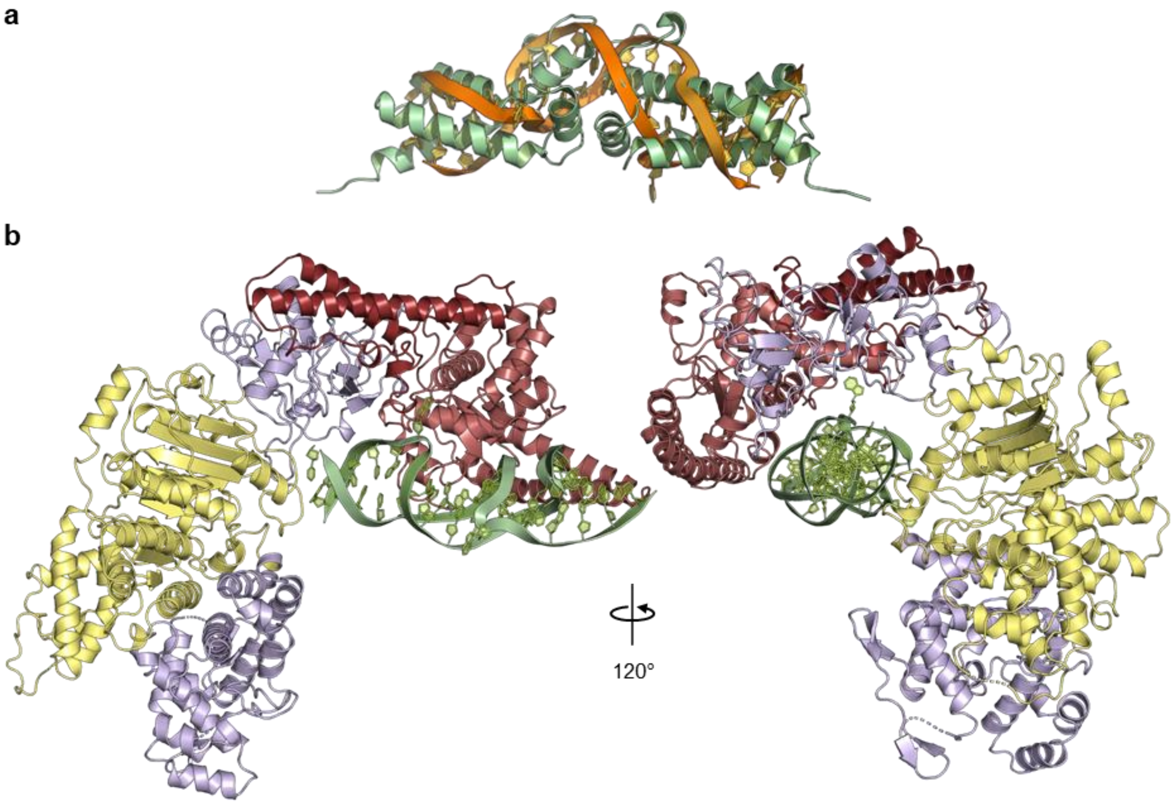


**Supplementary Figure S10. Ocr binding suggests that PglX binds DNA along the inside surface of the C-terminal domain.** (**a**) Alignment of an Ocr dimer (2Y7C) with its representative B-form DNA molecule (2Y7H). (**b**) Superposition of the DNA molecule from (a) onto the Ocr molecule in the PglX-SAM:Ocr complex.

**Supplementary Figure S11**

**a**

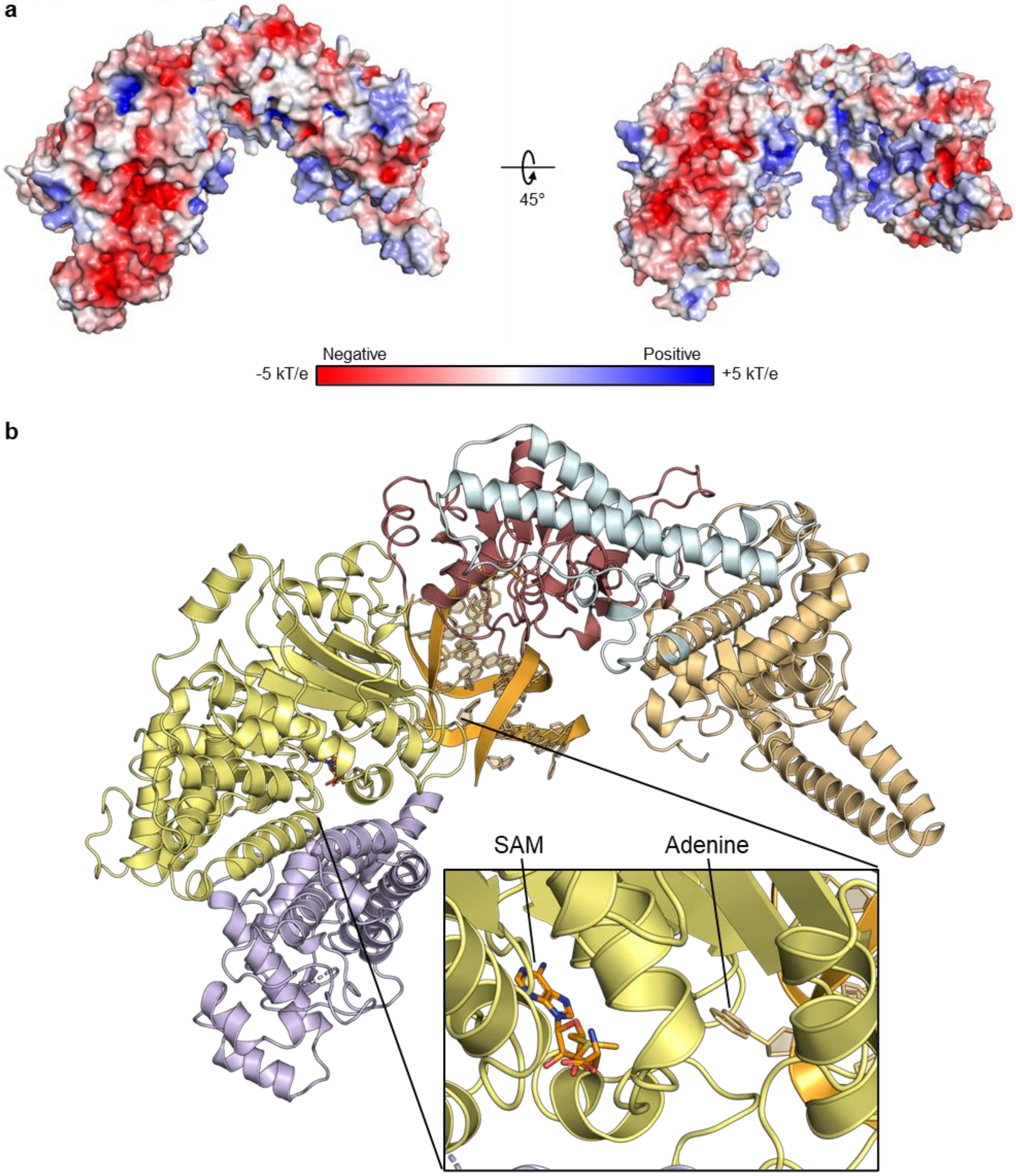


**Supplementary Figure S11. PglX displays a negatively charged surface within the predicted DNA binding region**. (**a**) Surface charge of PglX, showing a long negatively charged surface area along the C-terminal region into the target recognition domain. (**b**) Superposition of the 13 bp DNA molecule from the structure of MmeI (5HR4) onto the solved structure of PglX. Inset; the position of the flipped-out adenine base in the MmeI DNA molecule relative to the SAM molecule in PglX.

### Supplementary Figure S12

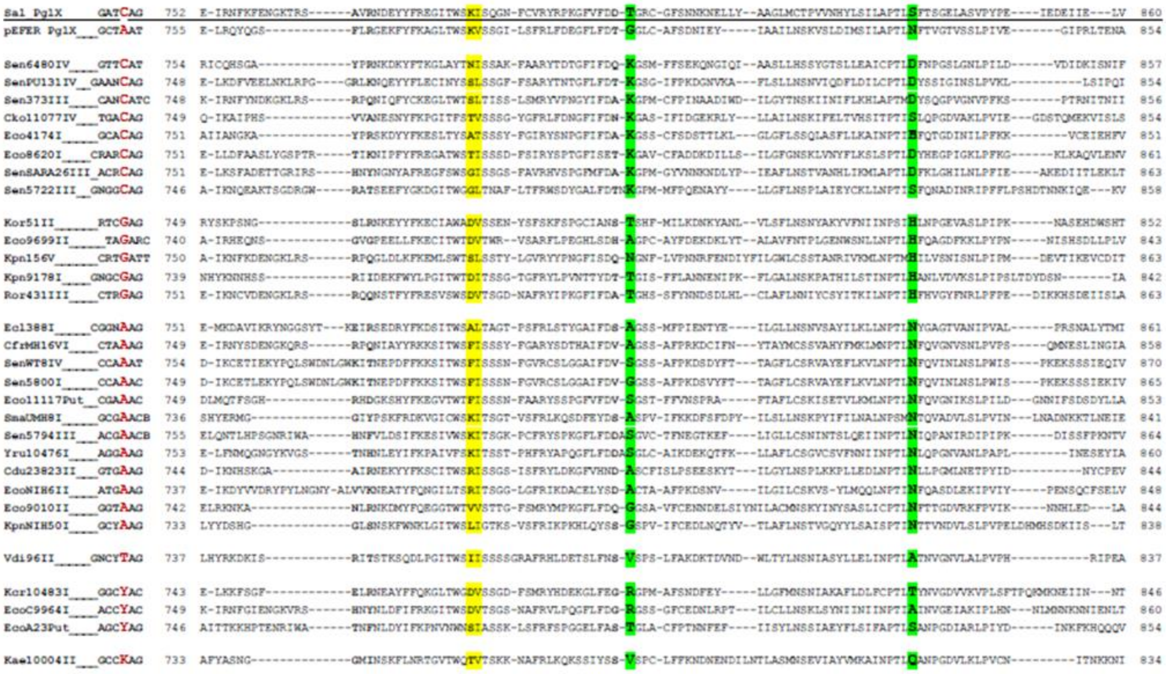

**Supplementary Figure S12. Example of the PglX alignments used for design of PglX mutants.** The residues in PglX which aligned with residues in MmeI controlling target sequence recognition were identified. The sequence of PglX was then aligned with homologues with known or predicted BREX recognition motifs and covariance between target residues and DNA recognition were identified. The above alignment represents identification of covariance of position -1 of the *Salmonella* BREX recognition motif (relative to the modified adenine) at PglX residues T802 and S838 (highlighted in green).

### Supplementary Tables

**Table S1. PglX mutants designed to alter DNA motif recognition.**

| Mutant number | Motif position[1] | Current motif base | Mutations | Predicted change | Resulting motif |
|---|---|---|---|---|---|
| 1 | -1 | C | T802T; S838H | G | GATGAG |
| 2 | | | T802N; S838H | G | GATGAG |
| 3 | | | T802A; S838N | A | GATAAG |
| 4 | | | T802G; S838N | A | GATAAG |
| 5 | | | T802V; S838A | T | GATTAG |
| 6 | -2 | T | K782F; D801V | A | GAACAG |
| 7 | | | K782R; D801D | G | GAGCAG |
| 8 | | | K782R; D801S | G | GAGCAG |
| 9 | | | K782D; D801S | C | GACCAG |
| 10 | | | K782A; D801S | N | GANCAG |
| 11 | -3 | A | A684L; S687R | C | GCTCAG |
| 12 | | | A684G; S687K | C | GCTCAG |
| 13 | | | A684K; S687D | G | GGTCAG |
| 14 | | | A684R; S687A | G | GGTCAG |
| 15 | | | A684H; S687S | G | GGTCAG |
| 16 | | | A684V; S687Q | T | GTTCAG |
| 17 | | | A684T; S687Q | T | GTTCAG |
| 18 | -4 | G | A766R; R768Q | C | CATCAG |
| 19 | | | A766R; R768D | C | CATCAG |
| 20 | | | A766H; R768F | A | AATCAG |
| 21 | | | A766T; R768H | A | AATCAG |
| 22 | | | A766V; R768A | T/N | T(N)ATCAG |
| 23 | +1 | G | Swap entire loop with pEFER PglX loop (AAs 591 - 600) | T | GATCAT |

[1] Relative to modified adenine base.

**Table S2. Bacterial strains and bacteriophages used in this study.**

| Bacterial Strains | | |
|---|---|---|
| **Strain** | **Genotype** | **Source** |
| *Escherichia coli* | | |
| DH5α | F- Φ80lacZΔM15 Δ(lacZYA-argF) U169 recA1 endA1 hsdR17 (rk-, mk+) phoA supE44 λ-thi1 gyrA96 relA1 | Invitrogen |
| ER2796 | K-12 F λ- fhuA2 Δ(lacZ)r1 glnV44 mcr-62 trp-31 dcm-6-zed-501::Tn10hisG1 argG6 rpsL104 dam-16::Kan xyl-7 mtlA2 metB1 (mcrB-hsd-mrr) 144::IS10 | NEB |
| BL21 (DE3) | B; $F^-$ *ompT* gal dcm lon $hsdS_B(r_B - m_B -)$ λ(DE3 [lacI lacUV5-T7p07 ind1 sam7 nin5]) $[malB^+]_{K-12}(\lambda^S)$ | Invitrogen |
| *Salmonella* D23850Δϕ | Δϕ(ΔBTP1 ΔBTP5 ΔGifsy-1 ΔGifsy-2 ΔST64B) | (Owen *et. al.*, 2017) |
| *Salmonella* D23850ΔϕΔBREX | Δϕ ΔBREX | (Rodwell *et al.*, 2021) |

| Bacteriophages | | | |
|---|---|---|---|
| **Phage** | **Host** | **Genbank accession** | **Source** |
| TB34 | *E. coli* DH5α | OX001802.1 | Environmental isolate (Kelly et. al., 2023) |
| Trib | *E. coli* DH5α | OX016465.1 | Environmental isolate (Kelly et. al., 2023) |
| Baz | *E. coli* DH5α | LR880803.1 | Environmental isolate (Kelly et. al., 2023) |
| Alma | *E. coli* DH5α | OV101294.1 | Environmental isolate (Kelly et. al., 2023) |
| Pau | *E. coli* DH5α | LR865361.1 | Environmental isolate (Kelly et. al., 2023) |
| BB1 | *E. coli* DH5α | MT843274.1 | Environmental isolate (Kelly et. al., 2023) |
| Jura | *E. coli* DH5α | LR999871.1 | Environmental isolate (Kelly et. al., 2023) |
| Mak | *E. coli* DH5α | OX001577.1 | Environmental isolate (Kelly et. al., 2023) |

| | | | |
|---|---|---|---|
| Bam | *E. coli* DH5α | OW991346.1 | Environmental isolate (Kelly et. al., 2023) |
| CS16 | *E. coli* DH5α | LR999870.1 | Environmental isolate (Kelly et. al., 2023) |
| Mav | *E. coli* DH5α | LR990702.1 | Environmental isolate (Kelly et. al., 2023) |
| Sip | *E. coli* DH5α | OU734268.1 | Environmental isolate (Kelly et. al., 2023) |
| T7 | *E. coli* DH5α | V01146.1 | Lab strain (ATCC BAA-1025-B2) |
| AWAQ | *Salmonella* D23850ΔϕΔBREX | Not sequenced | Environmental isolate, this study. |
| DA1 | *Salmonella* D23850ΔϕΔBREX | Not sequenced | Environmental isolate, this study. |
| DB1 | *Salmonella* D23850ΔϕΔBREX | Not sequenced | Environmental isolate, this study. |
| KMP | *Salmonella* D23850ΔϕΔBREX | Not sequenced | Environmental isolate, this study. |
| LTE | *Salmonella* D23850ΔϕΔBREX | Not sequenced | Environmental isolate, this study. |
| SB58 | *Salmonella* D23850ΔϕΔBREX | Not sequenced | Environmental isolate, this study. |
| SGP | *Salmonella* D23850ΔϕΔBREX | Not sequenced | Environmental isolate, this study. |
| SL2K | *Salmonella* D23850ΔϕΔBREX | Not sequenced | Environmental isolate, this study. |

**Table S3. Primers used in this study.**

| Primer | Sequence | Notes |
|---|---|---|
| **Gibson assembly primers for creation of pBrxXL$_{Sty}$ knockouts (KO)** | | |
| TRB1734 | CGATGAAAACGTTTCAGTTTGCTCATGGAAA ACGGTGTAA | pBrxXL$_{Sty}$ *pglX* Gibson assembly KO FWD (1) |
| TRB1735 | CGCAGGCGATCGTCGAAGCTTTACTGAAGAC GAATCCGGT | pBrxXL$_{Sty}$ *pglX* Gibson assembly KO REV (1) |
| TRB1736 | ACCGGATTCGTCTTCAGTAAAGCTTCGACGAT CGCCTGCG | pBrxXL$_{Sty}$ *pglX* Gibson assembly KO FWD (2) |
| TRB1737 | TTACACCGTTTTCCATGAGCAAACTGAAACGT TTTCATCGCTCTGGA | pBrxXL$_{Sty}$ *pglX* Gibson assembly REV (2) |
| TRB1738 | ATGCACCGGAGATTATTTAAATCTGGAATGA AATGGGATT | pBrxXL$_{Sty}$ PARIS Gibson assembly FWD (1) |
| TRB1739 | TTACACCGTTTTCCATGAGCAAACTGAAACGT TTTCATCG | pBrxXL$_{Sty}$ PARIS Gibson assembly REV (1) |
| TRB1740 | CGATGAAAACGTTTCAGTTTGCTCATGGAAA ACGGTGTAA | pBrxXL$_{Sty}$ PARIS Gibson assembly FWD (2) |
| TRB1741 | TTCTTCGTTGGTCAGAAACACGTATTTGTCTT CAACACGT | pBrxXL$_{Sty}$ PARIS Gibson assembly REV (2) |
| TRB1742 | ACGTGTTGAAGACAAATACGTGTTTCTGACC AACGAAGAA | pBrxXL$_{Sty}$ PARIS Gibson assembly FWD (3) |
| TRB1743 | AATCCCATTTCATTCCAGATTTAAATAATCTCC GGTGCAT | pBrxXL$_{Sty}$ PARIS Gibson assembly KO REV (3) |
| TRB1788 | GATGAAAACGTTTCAGTTTGCTCATGGAAAA CGGTGTAACA | pBrxXL$_{Sty}$ *brxC* Gibson assembly KO FWD (1) |
| TRB1790 | GAATCCGGTCACCTGCCTCAGCTGTGCTCTTT AACGAGGA | pBrxXL$_{Sty}$ *brxC* Gibson assembly KO REV (1) |
| TRB1791 | TCCTCGTTAAAGAGCACAGCTGAGGCAGGTG ACCGGATTC | pBrxXL$_{Sty}$ *brxC* Gibson assembly KO FWD (2) |
| TRB1792 | CCGGCGCGATCAGATAACTTTCAATACAAAA ACGCGGAAGAACC | pBrxXL$_{Sty}$ *brxC* Gibson assembly KO REV (2) |
| TRB1793 | CTTCCGCGTTTTTGTATTGAAAGTTATCTGAT CGCGCCGG | pBrxXL$_{Sty}$ *brxC* Gibson assembly KO FWD (3) |
| TRB1794 | GTTACACCGTTTTCCATGAGCAAACTGAAACG TTTTCATCGCT | pBrxXL$_{Sty}$ *brxC* Gibson assembly KO REV (3) |
| TRB1778 | GATGAAAACGTTTCAGTTTGCTCATGGAAAA CGGTGTAAC | pBrxXL$_{Sty}$ *ariA* Gibson assembly KO FWD (1) |
| TRB1795 | AGTGTTTTCAGTAAGTTGCCATCGAACTCATC AAGAATGC | pBrxXL$_{Sty}$ *ariA* Gibson assembly KO REV (1) |
| TRB1796 | GCATTCTTGATGAGTTCGATGGCAACTTACTG AAAACACT | pBrxXL$_{Sty}$ *ariA* Gibson assembly KO FWD (2) |
| TRB1797 | TTGCCATCGGTTAAATCAATTTAAATAATCTC CGGTGCAT | pBrxXL$_{Sty}$ *ariA* Gibson assembly KO REV (2) |
| TRB1798 | ATGCACCGGAGATTATTTAAATTGATTTAACC GATGGCAA | pBrxXL$_{Sty}$ *ariA* Gibson assembly KO FWD (3) |
| TRB1799 | GTTACACCGTTTTCCATGAGCAAACTGAAACG TTTTCATC | pBrxXL$_{Sty}$ *ariA* Gibson assembly KO REV (3) |

**Primers used for LIC cloning into pBAD30-LIC**

| | | |
|---|---|---|
| TRB1587 | caacagcagacgggaggtAATACCAATAACATCAAAAA | FWD LIC PglX Salmonella D23580 |
| TRB1588 | gcgagaaccaaggaaaggttattaAATAATCTCCGGTGCATTGC | REV LIC PglX Salmonella D23580 |
| TRB1856 | tggagccacccgcagttcgaaaaaTCAGGAGTCAAGACTGAG | FWD for cloning Strep site into pSAT1-LIC |
| TRB1857 | tttttcgaactgcgggtggctccaCGGATGATGATGATGATGATG | REV for cloning Strep site into pSAT1-LIC |
| TRB2025 | CAACAGCAGACGGGAGGTCCTCTAGAAATAATTTTGTTTAAC | FWD LIC pBAD30-RBS-His-Strep-PglX |
| TRB2026 | GCGAGAACCAAGGAAAGGTTATTAGTTATTAAATAATCTCCGGTG | REV LIC pBAD30-RBS-His-Strep-PglX |
| TRB2027 | CAACAGCAGACGGGAGGT GAAGGAGATATATCCATG AATACCAATAACATCAAAAAATAC | FWD LIC pBAD30-RBS-PglX |
| TRB2028 | GCGAGAACCAAGGAAAGGTTATTA TTATTAAATAATCTCCGGTGC | REV LIC pBAD30-RBS-PglX |

**Sequencing primers (ranges indicate position of primer in *Salmonella* BREX coding region)**

| | | |
|---|---|---|
| TRB710 | CGTTACCTGGAACCATTCGT | 2289-2308 |
| TRB711 | CCCTATGGATAGCTGGGATG | 3002-3021 |
| TRB712 | GCAGGACGTGATGGGTTTTA | 3688-3707 |
| TRB713 | GCCAATACGACGCGTTTAAG | 4403-4402 |
| TRB714 | GTCTATCCGGACCAAAGGTG | 5094-5113 |
| TRB715 | CGGCTGCATTTTAATTCGTT | 5796-5815 |
| TRB716 | GCACAAACTATGGCGGAAAT | 6499-6518 |
| TRB717 | ACGGATGCCGAGAAGAAGAT | 7195-7214 |
| TRB718 | GATAACCCGACAGGCTTTGA | 7888-7907 |
| TRB719 | GTCTCGACATTGACGACCG | 8600-8618 |
| TRB720 | ATTACGATGGCACATTTGGG | 9292-9311 |
| TRB721 | GGATTAATGTGCACTCCGGT | 10000-10020 |
| TRB722 | AAATCTCGAATTTATCGCCG | 10704-10723 |
| TRB723 | ATTGGCTGGGCACGGGTA | 11398-11415 |
| TRB724 | GGCGGTGTTGTACTCATTGAT | 12105-12126 |
| TRB725 | CCGGTTTTAATACTGCGTTTC | 12788-12808 |
| TRB726 | CTTGAAAGGCCTGGTCACTG | 13505-13524 |

| | | |
|---|---|---|
| TRB727 | TTGCTGAATCTGCGTAATCG | 14193-14212 |
| TRB728 | GCATAACACCATTGATGCCA | 14888-14907 |
| TRB729 | CGAATTTCAATCGCCGTAAT | 15598-15618 |
| TRB730 | GTATTTACCGCACGTACGCA | FOR_21 |
| TRB731 | AACCAGCGCGACGTTATC | FOR_22 |
| TRB732 | CGCGGTAGATATTCCGACTG | FOR_23 |
| TRB733 | CAACCTCATCCTCTTCACCTG | FOR_24 |

**Table S4. Plasmids used in this study.**

| Plasmid | Notes | Primers used | Reference |
|---|---|---|---|
| pBrxXL$_{Sty}$ | Full *S. enterica* serovar Typhimurium coding region in a pGGA vector backbone, created by golden gate assembly from genomic *S. enterica* serovar Typhimurium D23580 DNA. | TRB1367-TRB1378 | This study |
| pBrxXL$_{Sty}$-Δ*brxA* | Cloned by Genscript from pBrxXL$_{Sty}$. | Genscript synthesis | This study (Genscript) |
| pBrxXL$_{Sty}$-Δ*brxB* | Cloned by Genscript from pBrxXL$_{Sty}$. | Genscript synthesis | This study, (Genscript) |
| pBrxXL$_{Sty}$-Δ*brxC* | Created by Gibson assembly from pBrxXL$_{Sty}$. | TRB1788, TRB1790-TRB1794 | This study |
| pBrxXL$_{Sty}$-Δ*pglX* | Created by Gibson assembly from pBrxXL$_{Sty}$. | TRB1734-TRB1737 | This study |
| pBrxXL$_{Sty}$-Δ*pglZ* | Cloned by Genscript from pBrxXL$_{Sty}$. | TRB904/907 | This study, (Genscript) |
| pBrxXL$_{Sty}$-Δ*brxL* | Cloned by Genscript from pBrxXL$_{Sty}$. | TRB904/958 | This study |
| pBrxXL$_{Sty}$-Δ*ariA* | Created by Gibson assembly from pBrxXL$_{Sty}$. | TRB1778, TRB1795-TRB1799 | This study |
| pBrxXL$_{Sty}$-Δ*ariB* | Cloned by Genscript from pBrxXL$_{Sty}$. | Genscript synthesis | This study, (Genscript) |
| pBrxXL$_{Sty}$-Δ*ariA*Δ*ariB* | Created by Gibson assembly from pBrxXL$_{Sty}$. | TRB1748-TRB1743 | This study |
| pTRB507 | pGGA plasmid backbone containing 12400 – 14394 of the pEFER plasmid from *E. fergusonii,* used as a negative control. | | Picton *et al.* 2021 |
| pSAT1-LIC | pBAT4 derivative; pMB1 replicon | | Cai *et al.* 2020 |
| pSAT1-Strep-6xHis-SUMO | Created by GA | TRB1856 TRB1857 | This study |
| pSAT1-Strep-6xHis-SUMO-pglX | Created by LIC | TRB1587 TRB1588 | This study |
| pBAD30-Strep-6xHis-SUMO-pglx | Created by LIC | TRB2025 TRB2026 | This study |
| pBAD30-pglX | Created by GA | TRB2027 TRB2028 | This study |
| pBAD30-ocr | Created by LIC | TRB1911 TRB2081 | This study |
| pBAD30-*pglX*(mut.1–23) | Derived from pBAD30-*pglX.* Twenty-three plasmids each with different mutant *pglX* genes, described in Table S3. | Genscript synthesis | This study |
